# Insecticidal Efficacy of Selected Botanicals on three species of Adult Mosquitoes in Osogbo, Osun State, Nigeria

**DOI:** 10.1101/2023.02.11.528111

**Authors:** P.K. Ademodi, L.O. Busari, Z.O. Iwalewa, K.A Fasasi

## Abstract

The present study investigated the insecticidal potency of phytochemicals from three indigenous botanicals namely *Moringa oleifera* (Drumstick tree), *Vernonia amygdalina* (Bitter leaf) and *Ocimum gratissimum* (Scent leaf) against adult mosquitoes of the *Anopheles* species, *Aedes* species and *Culex* species cum their larval habitats.

Collection of the mosquito larval was made after scouting and identifying their breeding sites using scoops, plastic containers, sieves etc. Larva collection was done in the wet season (April to October) while the collected larva were conveyed to the laboratory and reared until they emerged into adults. A bioassay comprising twenty five adult mosquitoes from each of the three adult mosquito species were exposed to the botanicals at three different concentrations (5g/10ml, 10g/10ml and 15g/10ml) to assess their insecticidal potency which was measured through their knockdown rate (kdr) and mortality rate. The kdr were recorded at intervals for an hour while the mortality rates were recorded after 24hours of exposure.

A total of 400 mosquito larva were collected and five larval habitat encountered. The three botanicals showed insecticidal potency against the adult mosquito species. However, *O. gratissimum* had the highest potency at 15g/10ml against *Anopheles* (46%) and *Culex* (34%) species respectively (p □ 0.05). *Aedes spp* was resistant to the botanicals which may be due to a resistance barrier mechanism in *Aedes spp* that the botanicals lack the ability to break.

The botanicals could be adopted in the control of adult mosquitoes as they are biodegradable, eco-friendly and economical in contrast to synthetic insecticides with a view to consolidating global efforts on mosquito-borne diseases control and elimination with little or no human or environmental hazard.

## Introduction

Mosquitoes constitute a myriad of public health menace as vectors for most of the life-threatening diseases like malaria, yellow fever, dengue fever, chikungunya fever, filariasis, encephalitis, West Nile Virus infection, etc (Grigoraki *et al*., 2016). These infectious diseases mainly affect people in the tropics and thus regarded as tropical diseases (Rahuman *et al*., 2009). *Culex, Aedes* and *Anopheles* are the most important vectors involved in disease transmission to humans. About 229 million cases and 409 thousand deaths were reported due malaria incidence, while 56 million cases for dengue fever have been reported globally (WHO, 2020). Furthermore, more than ten million people were reported dead annually by other mosquito-borne diseases, such as filariasis, yellow fever, and encephalitis (WHO, 2021). There are 140 *Anopheles* species. 20 of these species are known to transmit malaria to human (Sinka *et al.*, 2012).

Resistance of mosquito vectors to synthetic insecticides globally is of great public health concern in vector control as it poses a great barrier towards global efforts on mosquito-borne diseases control and eradication. The cause of which has been linked to the indiscriminate use of synthetic insecticides (Grigoraki *et al*., 2016). Therefore, allowing for the easy escape of the mosquito vectors of the cidal effect of the insecticides through antigenic shifting and other resistance mechanisms. Thus, the need for the adoption of natural plant products (botanicals) which are usually non-toxic to man and domestic animals (Ejeta *et al*., 2021). These botanicals are also rich sources of bioactive organic chemicals and offer an advantage over synthetic insecticides as plants are less toxic, less prone to the development of resistance against them and easily biodegradable (Prabhu *et al*., 2011; Ajaegbu *et al*., 2016).

In Africa and other parts of the world, such as Ethiopia and China respectively, natural plants such as *O. gratissimum* and *Moringa oleifera* have been reported to be used as repellant against mosquito vectors (Prabhu *et al*., 2011; Ejeta *et al*., 2021). *Ocimum gratissimum* and *O. basilicum* have also been shown to have more repellant properties than plants such as *Azadirachta indica, Citrus lemon* and *Jatropha curcas* (Kazembe and Chaibva, 2012). Other botanicals whose insecticidal properties have been studied include *Calpurnia aurea, Momordica foetida, Zehneria scabra* etc (Muhammed *et al*., 2022).

Nigeria is reported to have the largest burden of NTDs in sub-Saharan Africa and accounting for a quarter (25%) of the total burden in the region (Hotez, 2009).13 out of the 20 NTDs are reported to be endemic in Nigeria (Hotez *et al*., 2012). Thus, the need for substantial work on mosquito-borne diseases prevention and control through vector control.

There is a paucity of reports in assessing the insecticidal efficacy of indigenous botanicals on mosquitoes in Osun State despite the state as well as other Southwestern states in the country endemicity for mosquito-borne diseases as reported by the WHO, 2021.

The present study therefore seeks to determine the efficacy of selected botanicals (through their phytochemical properties) on the selected adult mosquito species and their larva habitat in Osogbo, Osun state, southwest, Nigeria. This will assist in corroborating efforts targeted at mosquito-borne diseases control and elimination in the State and the world at large.

## Materials and Methods

The study area (Osogbo) is the capital of Osun State, Southwest, Nigeria. It covers an area of 14875 km^2^ with a population of 400000 (National Population census, 2016) and serves as a tourist centre due to the presence of the Osun groove. Agriculture is the primary occupation of residents and with a significant number of traders and artisans.

Ethical approval was obtained from the Osun State Ministry of Health, Abere, Osogbo, Osun State.

Thorough quest was made to identify the breeding sites of the mosquitoes. Larva collection was made using plastic containers, scoops, sieves etc. The collected larva was transported to the laboratory where they were bred till, they emerged into adults for laboratory bioassay.

The leaves of the botanicals used were collected from the trees within the town. These include *O. gratissimum* (scent leaf), *M. oleifera* (drumstick) and *Vernonia amygdalina* (bitter leaf).

We weighed 5g, 10g and 15g of each of the botanicals separately using weighing balance (Biobase electronic balance, model BE30001NF). Each mass was rinsed in a bowl of distilled water and then transferred into clean porcelain mortar and pounded using a pestle. About 10ml of acetone was added to each gram of the pounded botanicals to extract the plant material (phytochemical) which was sieved using a 0.55mm mesh sieve.

Thereafter extraction, the plant extracts were shake and impregnated on A4 papers by soaking them in the extracts. The impregnated papers containing the extracts are then allowed to dry for 10mins after which they are folded into WHO insecticide bioassay bottles. Four replicates comprising of 25 adult mosquitoes of each species was exposed to each bioassay bottles containing the plant extracts impregnated papers according to WHO standard by recording the knockdown rate (kdr) per hour and the mortality rate after 24hrs.

### Statistical Analysis

The data form the study was subjected to t-test and chi –square to test the significance in the efficacy among the botanicals.

## Results

Of the 5 mosquito larval habitats encountered during the study which included ground pools, gutters, tyres, containers and ponds. *Anopheles* and *Culex* larvae are both found breeding in ground pools and ponds while *Aedes* larva in tyres and containers. However, gutters were found as breeding sites for *Culex* larva alone (Table 1).

**Table 1:** Mosquito larva habitat during the study.

|  | Ground pool | Gutter | Tyres | Containers | Ponds |
| --- | --- | --- | --- | --- | --- |
| **Anopheles larvae** | + | — | — | — | + |
| **Culex larvae** | + | + | — | — | + |
| **Aedes larvae** | — | — | + | + | — |

Present: +; Absent: -

After 24 hours of exposure, a variation in the mortality of the female *Anopheles spp* was observed in the three botanicals used. At 15g/10ml concentration, the highest mortality was observed in *O. gratissimum* (46%), *V. amygdalina* (42%) and *M. oleifera* (30%) respectively (p=0.026; p<0.05). At 10g/10ml, the botanical that recorded the highest mortality was *M. oleifera* (44%), *O. gratissimum* (37%) and *V. amygdalina* (32%) respectively (p=0.024; p<0.05). Furthermore, the highest mortality at 5g/10ml was observed in *M. Oleifera* (38%), *O. gratissimum* (36%) and *V. amygdalina* (26%) respectively (p=0.033; p<0.05) (Figure 1).

**Figure 1:**
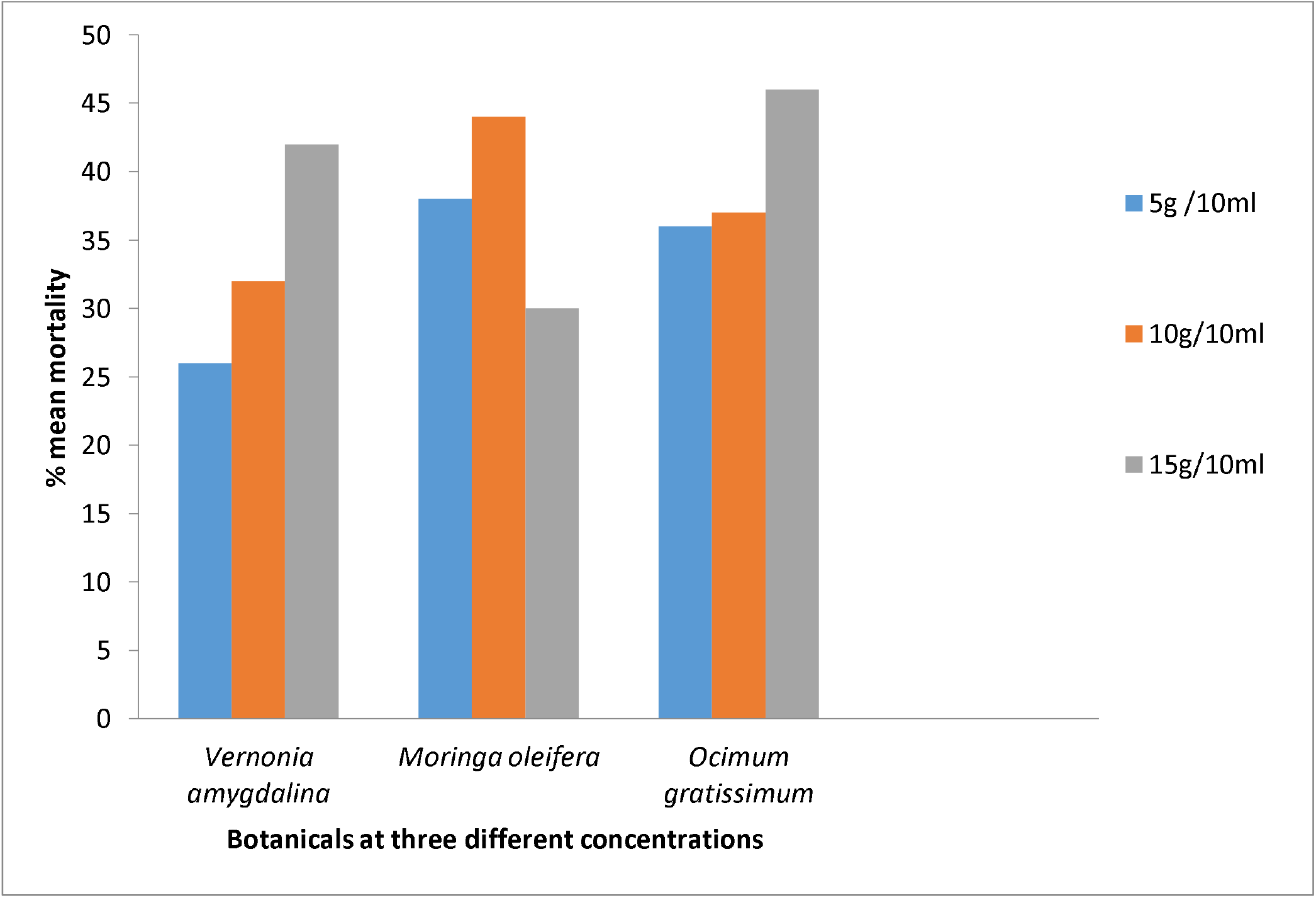
Comparative mortality of the botanicals on *Anopheles spp* at different concentrations during the study period.

There was a disparity in the mortality of the *Aedes spp* in the three botanicals used. At 15g/10ml concentration, the highest mortality was observed in *O. gratissimum* (42%), *M. oleifera* (11%) and *V. amygdalina* (4%) respectively (p=0.165; p>0.05). At 10g/10ml concentration, *M. oleifera* (17%) was observed to exhibit the highest mortality, *O. gratissimum* (11%) and *V. amygdalina* (7%) respectively (p=0.138; p>0.05). At 5g /10ml concentration, *O. gratissimum* (11%) had the highest mortality, *M. oleifera* (6%) and *V. amygdalina* (2%) respectively (p=0.183; p>0.05) (Figure 2).

**Figure 2:**
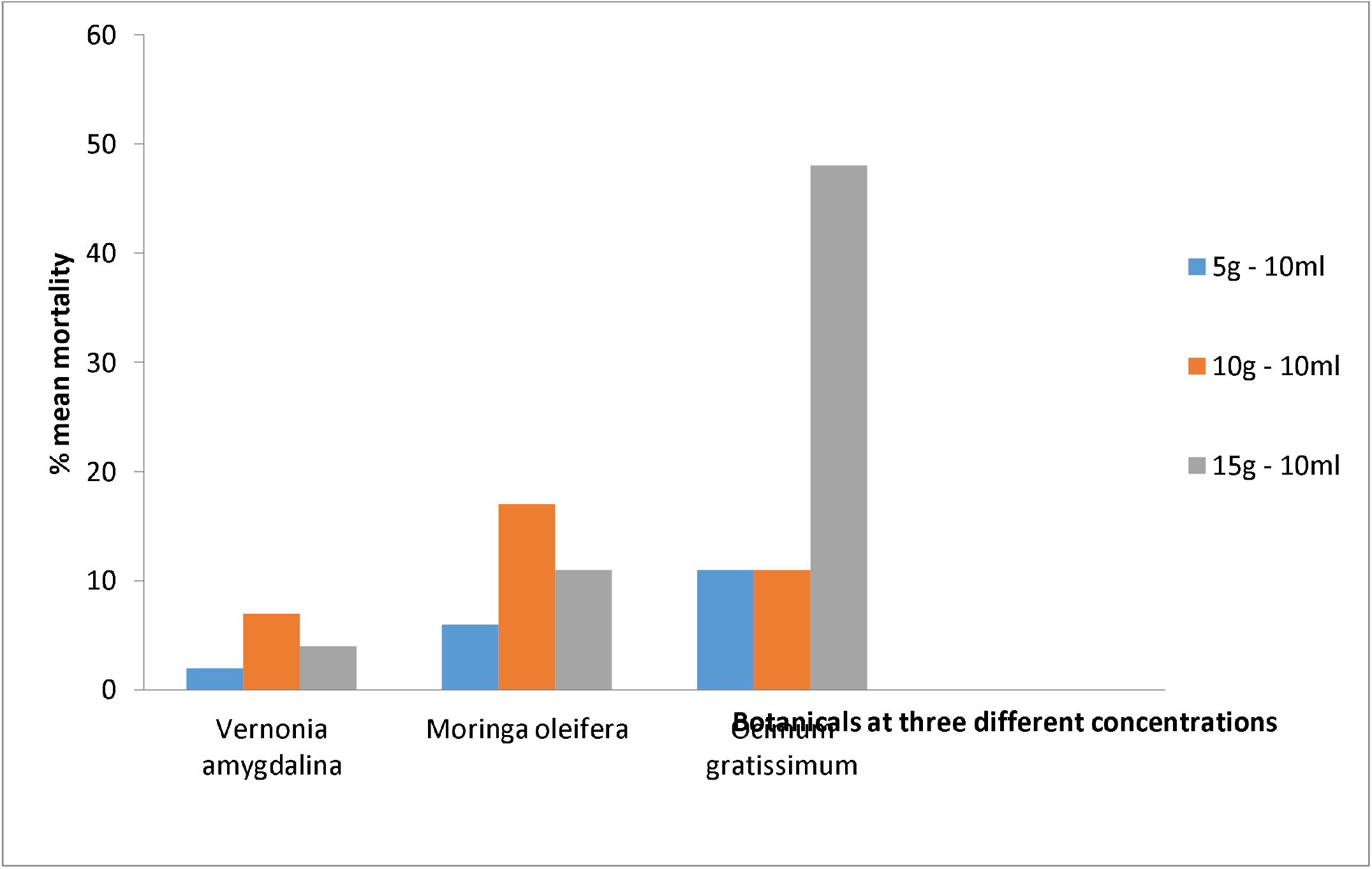
Comparative mortality of the botanicals on *Aedes* species at different concentrations during the study period.

After 24 hours of exposure, a variation in the mortality of the female *Culex* mosquito vectors was observed in the three botanicals used. At 15g/10ml concentration, *O. gratissimum* (34%) was recorded to have the highest mortality, then *V. amygdalina* (30%) and *M. oleifera* (20%) respectively (p=0.037; p<0.05). At 10g/10ml concentration, it was observed that *M. oleifera* (26%) had the highest mortality, *O. gratissimum* (23%) and *V. amygdalina* (13%) respectively (p=0.095; p>0.05). Lastly, at 5g /10ml concentration, the highest mortality was observed in *V. amygdalina* (24%), *O. gratissimum* (22%) and *M. oleifera* (21%) respectively (p=0.004; p<0.05) (Figure 3).

**Fig 3:**
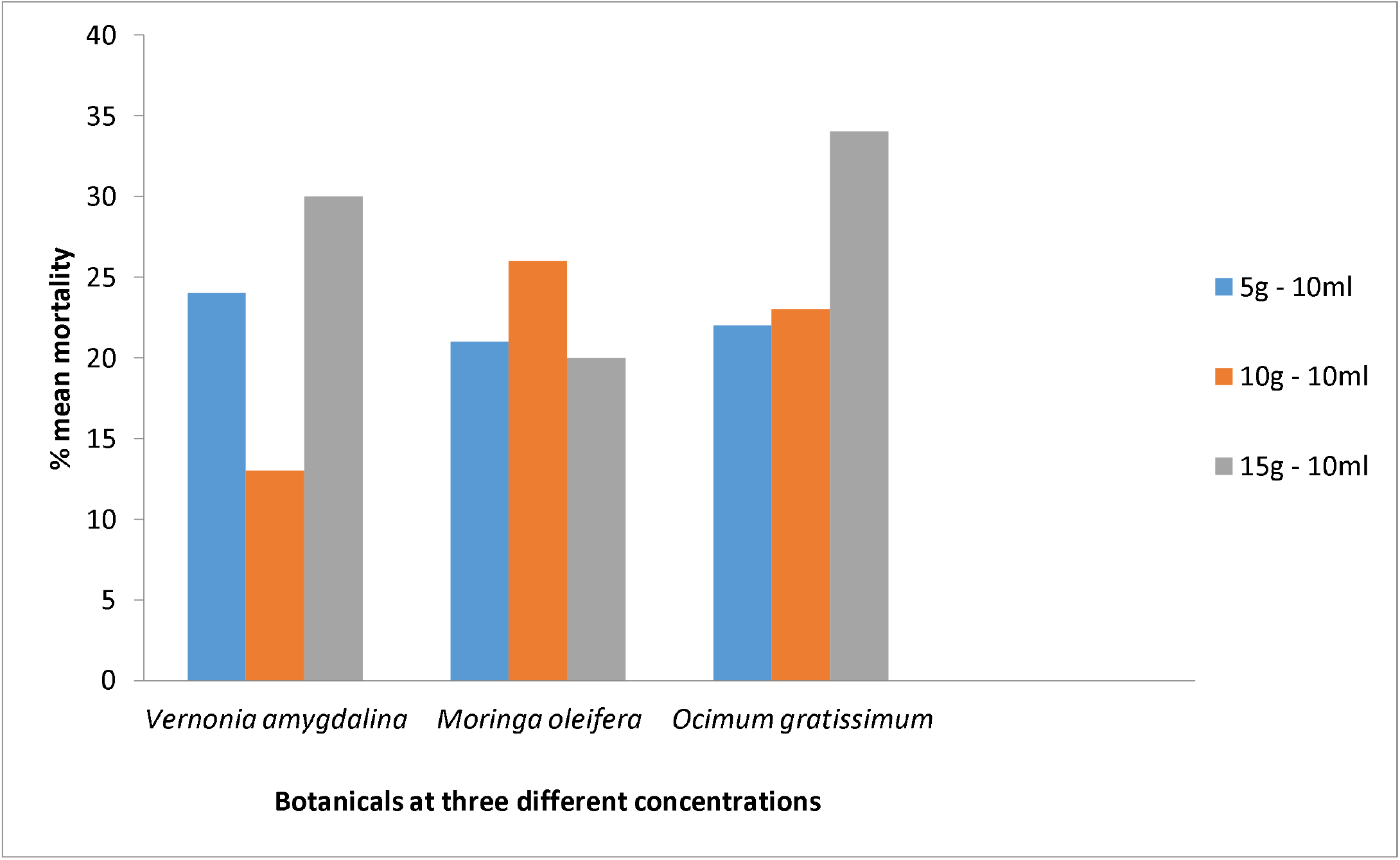
Comparative mortality effect of the botanicals on *Culex* species at different concentrations during the study period.

At 5g/10ml, the highest susceptibility (38%) was observed in *Anopheles spp* with *M. oleifera* while the lowest (2%) in *Aedes spp* with *V. amygdalina* (p=0.001; p<0.05). *M. oleifera* had the highest mortality (44%) at 10g/10ml against *Anopheles spp* with the least (7%) in *Aedes spp* with *V*.*amygdalina* (p=0.005; p<0.05). However, at 15g/10ml, *O. gratissimum* gave the highest mortality (48%) against *Aedes spp* while the lowest (4%) was recorded in *Aedes spp* with *V. amygdalina* (p=0.38; p>0.05) (Table 2).

**Table 2:**
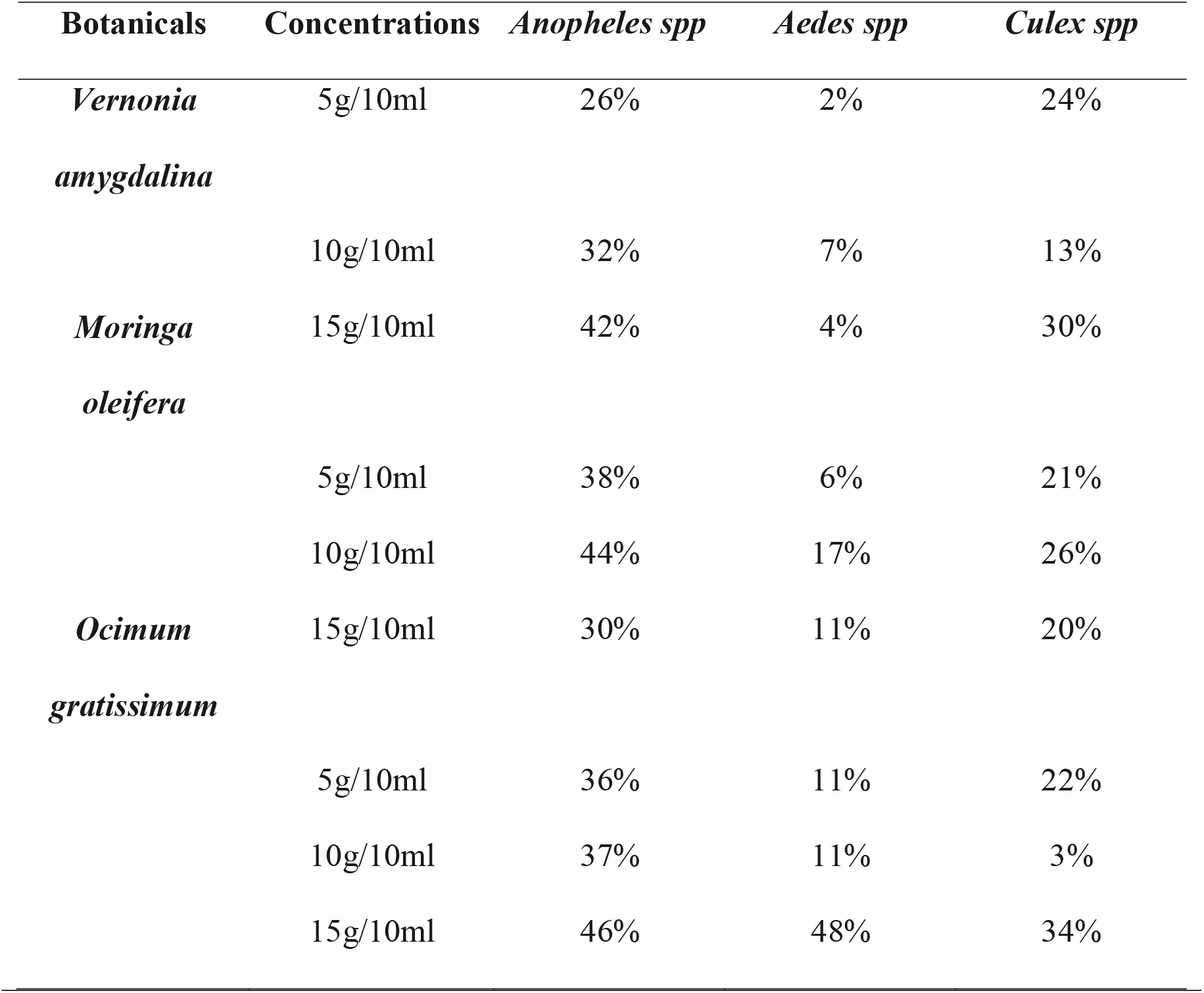
Susceptibility of the female mosquitoes to the three botanicals

At 5g/10ml the highest resistance (72%) was observed in *Aedes spp* with *V. amygdalina* while the lowest resistance (22%) was observed in *Anopheles spp* with both *V*.*amygdalina* and *M. oleifera* (p=0.001; p < 0.05). *V. amygdalina* had the highest resistance (64%) at 10g/10ml against *Aedes spp* while the least (17%) was recorded in *Anopheles spp* with *O. gratissimum* (p=0.021; p < 0.05). lastly, at 15g/10ml the highest resistance (77%) was observed in *Aedes spp* with *V. amygdalina* while the least (1%) was recorded in *Anopheles spp* with *O. gratissimum* (p=0,066; p > 0.05) (Table 3).

**Table 3:** Resistance of the female mosquitoes to the three botanicals

| **Botanicals** | **Concentration** | *Anopheles spp* | *Aedes spp* | *Culex spp* |
| --- | --- | --- | --- | --- |
| *Vernonia amygdalina* | 5g/10ml | 22% | 72% | 48% |
|  | 10g/10ml | 23% | 64% | 46% |
|  | 15g/10ml | 26% | 77% | 45% |
| *Moringa oleifera* | 5g/10ml | 22% | 54% | 42% |
|  | 10g/10ml | 25% | 55% | 20% |
|  | 15g/10ml | 23% | 41% | 48% |
| *Ocimum gratissimum* | 5g/10ml | 27% | 70% | 56% |
|  | 10g/10ml | 17% | 50% | 51% |
|  | 15g/10ml | 1% | 36% | 42% |

## Discussion

The scourge of emergence of mosquito resistance to synthetic insecticides is considered the main reason necessitating the need to look inward for new sustainable control measures. Plant extracts fulfilled such concern through their mild impacts on the non-target organisms (Amina *et al*., 2019). *Anopheles spp* were susceptible to the botanicals at the different concentrations. *Vernonia amygdalina, Moringa oleifera* and *Ocimum gratissimum* are insecticidal on adult *Anopheles spp* with *O. gratissimum* (46%) having the highest mortality (5) effect (p=0.026; p<0.05) at 15g/10ml. However, at 10g/10ml and 5g/10ml, *M. oleifera* had the highest mortality (44%) (The percentage may be better in figure) effect on the *Anopheles spp*. This study is in consonance with the observation of (Etta et al., 2016) who also reported the susceptibility of *Anopheles spp* to *O. gratissimum* plant extract. This also conforms to the study of Ashfaq and Ashfaq (2012) who reported that seed extract of *M. oleifera* exhibited excellent larvicidal, pupicidal and repelling action against *Culex spp* and can effectively be used to end the awe of dengue fever vectors. *Ocimum gratissimum* contains rich array of phytochemicals and essential oils which may be implicated in the insecticidal ability of its extract against mosquitoes (Etta et al., 2016). Okorie *et al*., (2020) reported that the presence of alkaloids, tannins, saponins, flavonoids and glycosides contributed to the insecticidal properties of the leaves.

*Culex spp* were only susceptible at 5g/10ml and 15g/10ml concentration respectively to the botanicals (p<0.05). At 5g/10ml and 15g/10ml concentrations, the highest mortality (percentage in figure) was observed in *Culex spp* with *V. amygdalina* (24%) and *O. gratissimum* (34%) respectively (p=0.004;p=0.037;p<0.05). It conforms to the study of Awosolu *et al*., (2018) whose result showed that the extract of *O. gratissimum* used as a repellant on *Culex* mosquitoes ‘population is effective. The present study shows that *Aedes spp* were not susceptible to the botanicals (p>0.05), the reason for which may be due to the inability of the phytochemicals from the botanicals to culminate the resistant mechanism in *Aedes spp*.

## Conclusion

The present study has documented the insecticidal efficacy of *O. gratissimum* as the most effective for *Anopheles spp*. among the three botanicals used. This would assist in vector control programmes targeted at malaria prevention and elimination both locally and internationally due to the public health threat posed by mosquito-borne diseases globally. However, further study on the resistance of *Aedes spp* to the botanicals is required since they are vectors of life-threatening diseases such as dengue fever etc. with a view to controlling and eradicating them.

